# Genetically-Modified Macrophages Accelerate Myelin Repair

**DOI:** 10.1101/2020.10.28.358705

**Authors:** Marie-Stephane Aigrot, Clara Barthelemy, Sarah Moyon, Gaelle Dufayet-Chaffaud, Leire Izagirre-Urizar, Beatrix Gillet-Legrand, Laura Bayón-Cordero, Satoru Tada, Juan-Carlos Chara, Carlos Matute, Nathalie Cartier, Catherine Lubetzki, Vanja Tepavčević

**Author notes:** Corresponding authors: VT, NC, CL. Note-current address for Satoru Tada: Department of Neurology, Osaka University Graduate School of Medicine, 2-2 Yamadaoka, Suita, Osaka 565-0871, JAPAN.

## Abstract

Preventing neurodegeneration-associated disability progression in patients with multiple sclerosis (MS) remains an unmet therapeutic need. As remyelination prevents degeneration of demyelinated axons, promoting this process in patients might halt the development of permanent disability. In demyelinating mouse lesions, local overexpression of Semaphorin 3F (Sema3F), an oligodendrocyte progenitor cell (OPC) attractant, increases OPC recruitment and remyelination. However, molecular targeting to MS lesions is a challenge because these are disseminated in the central nervous system.

We hypothesized that a clinically-relevant paradigm to deliver Sema3F to demyelinating lesions and increase OPC recruitment may be to use blood-derived macrophages as vehicles. Thus, we chose transplantation of genetically-modified hematopoietic stem cells (HSCs) as means of obtaining circulating monocytes that overexpress Sema3F. We first demonstrated that the supernatant from Sema3F-lentiviral vector transduced HSCs stimulates OPC migration in Neuropilin 2 (Nrp2, Sema3F receptor)-dependent fashion. We then investigated whether OPCs remain responsive to Sema3F with age. While Sema3F expression in the lesions of middle-aged and old mice (characterized by decreased efficiency of OPC recruitment and remyelination) was decreased, middle-aged OPCs retained Nrp2 expression and migrated in response to both recombinant Sema3F and Sema3F-transduced cell supernatant in vitro. We then investigated whether blood cells engineered to overexpress Sema3F can target demyelinating CNS lesions and improve remyelination. Thus, we transplanted Sema3F-transduced HSCs and obtained chimeric mice (with Sema3F overexpression in blood cells), in which we induced demyelinating spinal cord lesions. Transgene-carrying cells, predominantly macrophages, quickly infiltrated lesions in both control and Sema3F chimeras. While infiltration of Sema3F-expressing cells did not alter the inflammatory status of the lesions nor OPC survival, it increased OPC recruitment, which accelerated the onset of remyelination. Our results provide a proof-of-concept that blood cells, particularly monocyte-derived macrophages, can be used to deliver pro-remyelinating agents “at the right time and place”, suggesting novel means for remyelination-promoting strategies.

## INTRODUCTION

Halting the progression of neurological disability in patients with multiple sclerosis (MS) is a major challenge. MS frequently starts as a relapsing-remitting disease (RRMS), with neurological symptoms alternating with episodes of recovery. RRMS eventually evolves into progressive disease, characterized by development of permanent neurological handicap. A subset of patients directly enters the progressive phase^1^.

Currently used treatments for MS are mostly immunomodulators and immunosuppressants. These reduce inflammation and related relapses, but are largely ineffective in preventing transition into progressive disease^1^ where accumulation of disability is the consequence of neuronal/axonal loss, likely triggered by increased vulnerability of demyelinated axons^2^. Remyelination (myelin regeneration) of demyelinated axons protects these from degeneration, as shown by neuropathological studies of MS tissue^2^, experiments using a model of demyelination^3^, and myelin PET imaging of MS patients^4^. Hence, developing remyelination-promoting strategies is a major therapeutic goal to prevent progression in patients with MS^5^.

In a subset of MS lesions, remyelination failure is associated with depletion/low numbers of oligodendroglial cells^6–9^. In these, stimulating oligodendrocyte progenitor cell (OPC) repopulation/recruitment to demyelinated areas appears crucial to enhance remyelination and achieve neuroprotection.

OPC recruitment in experimental lesions can be achieved by locally injecting a viral vector overexpressing a guidance molecule, Semaphorin 3F (Sema3F), an OPC attractant^8,10^. Moreover, we previously showed that OPCs in MS lesions express Sema3F receptor, Neuropilin 2 (Nrp2), and that while Sema3F is detected in early lesions, it is not expressed in chronically demyelinated lesions^11^. Thus, these data suggest that OPC recruitment in MS lesions may be enhanced by increasing intra-lesional Sema3F expression. However, the viral vector injection paradigm is not applicable in the patient setting for a number of reasons, one of them being that MS lesions are disseminated within the CNS. This question of how to optimally deliver therapeutic molecules to demyelinating MS lesions applies not only to strategies to increase OPC recruitment but also to other neuroprotective approaches such as delivering neurotrophic or antioxidative molecules to protect the axons, and in general, to any attempts to modify intralesional events in order to stop the progression of the pathology.

In leukodystrophies, a group of diseases in which a molecular defect in glial cells leads to anomalies in white matter formation and/or maintenance, intraparenchymal delivery of therapeutic proteins has been achieved using transplantation of autologous genetically-modified hematopoietic stem/progenitor cells (HSCs). Using this strategy, blood-born cells engineered to overexpress the missing brain protein were reported to infiltrate the brain, which raised parenchymal levels of the protein in question and corrected, at least in part, the associated myelin pathology. This approach has been successful in both mouse models and human leukodystrophy patients^12–17^.

We hypothesized that intralesional targeting of therapeutic molecules using blood-born cells could be a clinically-relevant strategy to stimulate remyelination in demyelinating CNS diseases, such as MS, by means of targeting the expression of Sema3F to demyelinating lesions and increasing OPC recruitment.

We first showed that monocytes/macrophages derived from transplanted HSCs efficiently infiltrate demyelinating mouse lesions. We then designed lentiviral vectors that allowed us to genetically modify HSCs to overexpress Sema3F and investigated whether these cells can stimulate OPC migration in vitro, as well as OPC recruitment to demyelinating lesions and remyelination in vivo. We showed that the supernatant of Sema3F-transduced HSCs increases migration of young and middle-aged OPCs in vitro in Nrp2-dependent manner, and that chimeric mice with Sema3F-overexpressing blood cells show increased OPC recruitment and accelerated onset of remyelination. Thus, we conclude that genetically-modified blood cells, particularly macrophages, can be used to efficiently and quickly express pro-remyelinating molecules in demyelinating lesions and promptly stimulate myelin repair.

Our results represent a proof-of-concept that genetically-modified monocytes/macrophages can be used to enhance CNS remyelination, which provides novel cues for therapies to prevent/diminish neurological disability in patients with MS.

## METHODS

### Mice

Five to 7 week-old female and 9-week old male C57/BL6 mice were purchased from Janvier (France). All animal manipulations were performed after a period of adaptation to the animal house of at least one week. The experiments were performed according to the European Union regulations and approved by the ethical committee for animal use (approval number 18471). Mice were housed under standard conditions and given ad libitum access to dry food and water.

### Isolation and culture of mouse hematopoietic stem/progenitor cells (HSCs)

Six-8 week-old female mice were sacrificed by cervical dislocation. Tibias and femurs were dissected and bone marrow flushed using a syringe. Differentiated white blood cells were then removed using Direct Lineage Cell Depletion Kit (Miltenyi Biotec). The resulting mixture of hematopoietic stem and progenitor cells was then kept in culture in IMDM (Sigma) supplemented with FBS (Sigma), and mSCF, hIL6, and mIL3 (all from Peprotech) in 96-well plates for 16-24 hours at 37°C.

### Lentiviral vector generation

DNA sequence encoding green fluorescent protein (GFP) alone or GFP and human Semaphorin 3F^18^, separated by a self-cleaving T2A sequence, were synthesised and inserted into the self-inactivated vector (SIN) backbone containing the WPRE element and the murine ubiquitous phosphoglycerate kinase 1 promoter (PGK). Lentiviral vectors were generated at the Molecular biology and vector production platform, MIRCEN, François Jacob Institute of Biology, CEA, (Fontenay-aux-Roses, France), as described previously^19^. The SIN vectors were pseudotyped with VSVg glycoprotein G. Viral particles were produced in human embryonic kidney (HEK)-293 T cells by a four-plasmid transient transfection system. The supernatant was collected 48 h later and filtered, and the particle content of the viral batches was determined by ELISA for the p24 antigen (Gentaur, Paris, France). High-titre stocks were obtained by ultracentrifugation. The pellet was re-suspended in 1% BSA in PBS, frozen and stored at −80 °C.

### Transduction of HEK cells and supernatant collection for migration assays

HEK 293T cells were grown in DMEM-Glutamax™ medium (Gibco) supplemented with 10% foetal calf serum at 37°C and 5% CO_2_. Eighty per cent confluent cells were transduced with PGK-GFP-T2A-Sema3F or PGK-GFP lentiviral vectors at a Multiplicity of Infection (MOI) of 10-20 for 24h. Medium was replaced and transduced cells were grown for 3-5 days. Supernatant was collected and stored at −80 °C.

### Lentiviral transduction of HSCs

Lentiviral transductions were performed in IMDM medium supplemented with FBS, mSCF, hIL6, mIL3, and protamine sulfate (Sigma). PGK-GFP and PGK-GFP-T2A-Semaphorin 3F vectors were applied at an MOI of 25, and cells were incubated at 37°C for 24 hours. The next day, medium was replaced with the fresh vector-containing medium and another round of transduction was performed. Control cells were incubated without lentiviral vectors. For supernatant collection, medium was replaced and transduced cells were grown for 3-5 days. Supernatant was collected and stored at −80 °C.

### In vitro expansion, transduction efficiency and viability test

Transduced cells were collected, washed, counted, and resuspended in IMDM medium with FBS (Sigma), and mSCF, hIL6, and mIL3 (all from Peprotech) in 6-well plates at initial density of 10^5^ cells/mL. Every other day, fresh medium was added to the well, and cells were passaged to new wells as needed. After 5 days of culture, cells were collected, washed, counted and resuspended in PBS. GFP expression (endogenous fluorescence) and viability (propidium iodide (PI) and Viobility 405/520 Fixable Dye (Miltenyi Biotec) staining) were investigated using flow cytometry (LSR Fortressa, Benton Dickinson).

### Colony-forming unit assay

Transduced cells were collected, washed, and resuspended in MethoCult™ GF M3434 medium (Stemcell) at a density of 1000 cells/mL and grown in 6-well plates (1000 cells/well) for 10-11 days. Numbers of erythroid, myeloid, and GFP+ colonies were counted and compared between the conditions.

### Western blot

Following transduction and 5-day expansion, cells were pelleted by centrifugation, and the supernatants were collected. Cell pellets were lysed in a lysis buffer containing protease inhibitor cocktail (Sigma or Thermofisher). Both the protein extracts from cell pellets and the supernatants from cultured cells were subjected to protein quantification using BCA Protein Assay Kit (Thermofisher). Samples were boiled at 99 °C for 2 min, size-separated by SDS-PAGE in 4–20% Criterion TGX Precast gels and transferred using Trans-Blot Turbo Midi Nitrocellulose Transfer Packs (Bio-Rad). Membranes were blocked with 5% BSA in Tris-buffered saline/0.5 % Tween-20 (TBS-T) for 1h at RT. Primary antibodies (Goat anti-GFP biotin conjugated, Vector Labs; sheep polyclonal anti-human-sema3f antibody, R&D systems; mouse anti-GADPH, Abcam) were diluted in the blocking buffer, and the membranes were incubated with antibody solution under gentle shaking overnight at 4 °C. Following 3 washes in TBS/Tween-20 0.1%, Biotin-conjugated polyclonal anti-sheep IgG (Vector Labs) in the case of anti Sema3F antibody was applied for 1 h at RT. HRP-conjugated streptavidin (Vector Labs) was applied for 1 h. For anti GADPH labelling, HRP-conjugated Donkey anti-Mouse IgG (Thermofisher) was applied. Bands were visualized using enhanced chemiluminescence detection kit (BioRad). Images were acquired using a Fusion FX imaging chamber (Vilber).

### In vitro migration assay

Purified OPCs, isolated using FACS from the brains of 2 month-old (young) and 9 month-old (middle-aged) PDGFRα-GFP mice, as previously described^6,10^, were resuspended in a modified Bottenstein-Sato (BS) medium and plated at the upper face of the porous membrane of the transwell chamber (Corning Costar Co, USA) positioned over the well within the 24-well plate (Corning) at a density of 10000 cells per membrane. Five hundred microliters containing a 1:1 mixture of the BS medium and supernatant from either non-transduced (NT)-, PGK-GFP transduced-, or PGK-GFP-T2A-Sema3F-transduced HSC or HEK cells were added to the well. Anti-Nrp2 function blocking antibody (AF2215, R&D Systems) was applied at a concentration of 10 µg/mL. Following a 24-hour incubation period, the cells present at the upper surface of the membrane were scraped off and the membrane fixed using 4% paraformaldehyde (PFA) solution. Nuclei were labelled with Hoechst and cells present on the lower surface of the membrane counted. The assays were performed as duplicates or triplicates. The statistical analyses were performed on replicates normalized to the average number of migrated control cells from the same experiment from 3-5 independent experiments and presented as fold change in the number of cells crossing the membrane insert relative to the control (NT) for each independent experiment.

### Preconditioning and cell transplantation

Recipient mice (10-12 week-old males) were injected (intraperitoneally) with cyclophosphamide (Sigma) at 200 mg/kg for 2 consecutive days (total of 400 mg/kg), and then with busulfan (Sigma) at 25 mg/kg for 4 consecutive days (total of 100 mg/kg). This procedure inhibits the proliferation of recipient cells and enhances transplanted cell integration.

The day following the last injection of busulfan, transduced cells were collected, washed, and resuspended in alphaMEM (Fisher Scientific) at a density of 3000 cells/µ L. Pre-conditioned mice were anaesthetized with isofluorane, and 200µ L of cell suspension (600000 cells total) was injected retro-orbitally. Mice were randomly selected to receive GFP- or Sema3F-GFP-transduced cell preparation. Mice were treated with Baytril for 1 month following preconditioning and transplantation, and their weight was recorded weekly.

### Detection of blood chimerism for transplanted cells

Presence of donor cells was investigated in the blood of transplanted mice using qPCR for sex determining region of the chromosome Y (SRY). Briefly, at 8 weeks post-transplantation, blood was isolated from transplanted mice by submandibular bleed, and gDNA was extracted. qPCR was performed using primers specific for actin (Sigma) and SRY (Thermofisher). Quantification was performed using the 2^-ΔΔCt^ algorithm.

### Detection and characterization of transduced cells in the blood of transplanted mice

To detect presence of cells transduced with lentiviral vectors in the blood of transplanted mice, blood was isolated at 8 weeks following transplantation. Red blood cells were lysed using a Red Blood Cell Lysis Buffer (Thermofisher), the cells were washed and the proportion of GFP+ cells was investigated using flow cytometry.

To characterize transduced cells in the blood, labeling with antibodies specific for different hematopoietic lineages was performed: CD11b (BD Bioscience; monocyte marker), CD3 (BD Bioscience; T cell marker), and CD19 (Bd Bioscience; B cells marker). The viability was investigated using Viobility 405/520 Fixable Dye (Life Technologies). All markers were applied at 1:10 dilution. Cells were analyzed using the cytometer LSR Fortessa (Benton Dickinson).

### Induction of demyelinating lesions

At 10 weeks post HSC transplantation, demyelinating lesions were induced in the spinal cord by a stereotaxic injection of 0.5µ L of 1% lysophosphatidylcholine (LPC, Sigma Aldrich) in sterile 0.9% NaCl solution. Chimeric mice were at least 20 weeks old at the time of lesion induction. Prior to the surgery mice were anaesthetized by intraperitoneal injection of ketamine (90 mg/kg) and xylazine (20 mg/kg) cocktail. Two longitudinal incisions into longissimus dorsi at each side of the vertebral column were performed and the muscle tissue covering the column was removed. Animals were placed in a stereotaxic frame, the 13th thoracic vertebra fixed in between the bars designed for manipulations on mouse spinal cord (Stoelting, USA) and intravertebral space exposed by removing the connective tissue. An incision into dura mater was performed using a 30G needle and LPC injected using a glass micropipette attached via a connector to Hamilton’s syringe and mounted on a stereotaxic micromanipulator. The lesion site was marked with sterile charcoal. Following LPC injection, the muscle sheaths were sutured with 3/0 monocryl, and the skin incision closed with 4/0 silk.

### Perfusion and tissue processing

Mice were euthanized with Imalgene and perfused with a 4% PFA (Sigma) solution in phosphate buffered saline (PBS; pH 7.4) for immunohistochemical (IHC) analysis or 4% glutaraldehyde (Electron Microscopy Sciences) solution supplemented with calcium chloride in 0.1 M phosphate buffer (PB) for transmission electron microscopy (TEM). For IHC analysis, spinal cords were re-equilibrated in 15% sucrose solution and frozen in 15% sucrose-7% gelatine solution in PBS. Twelve-micron coronal sections were cut using cryostat (Leica). For EM analyses, spinal cords were post-fixed overnight, washed in 0.1M PB and cut into four 1-mm-thick transverse blocks surrounding the injection area. The next day, the tissue was post-fixed in 1% osmium for 2 h, rinsed, dehydrated, and embedded in Epon-812 resin (EMS).

### Antibodies and immunohistochemistry

Primary antibodies used for immunohistochemistry were: Millipore antibodies: chicken anti-MBP (AB9348, 1:200), mouse anti-Olig2 (MABN50; 1:200), rabbit anti-GFP (AB3080, 1:500), mouse anti-MOG (MAB5680, 1:200); BD Biosciences antibodies: rat anti-PDGFRα (558774, 1:200), and rat anti CD45R/B220 (557390, 1:200); rabbit anti-cleaved caspase 3 (9661, Cell Signalling, 1:400); mouse anti-APC (OP80, Calbiochem; 1:200); guinea pig anti-Iba1 (Synaptic Systems; 234004, 1:300); Serotec antibodies: rat anti-CD68 (MCA19575, 1:200) rat anti-CD11b (MCA711; 1:200), and rat-anti CD3 (MCA1477, 1:50); rabbit anti-Iba1 (Wako; 019-19741; 1:250); rat anti-Sema3F (R&D Systems, MAB3237, 1:50); rabbit anti-P2Y12 (Anaspec, AS-55043A, 1:200). Secondary antibodies used were Alexa-conjugated goat antibodies (Invitrogen) and were used at 1:1000.

Sections were air-dried for 1h and re-hydrated in TBS. For all stainings but that for PDGFRα, antigen retrieval was performed by heating the sections in a low pH retrieval buffer (Vector Labs) at high power using a microwave for 45 seconds. After washing in TBS, slides were incubated in absolute ethanol solution for 15 min at −20°C. After washing, blocking buffer (TBS, 5% NGS, 1%BSA, 0.1% Triton or BlockAid solution (Thermofisher)) was applied for 30 min, followed by primary antibody incubation overnight at 4°C. In the case of PDGFRα, Triton was not used for overnight incubation. Sections were washed and secondary antibodies applied for 1 h. After washing, nuclei were counterstained with DAPI. Slides were mounted in Fluoromount-G (Thermofisher Scientific).

### Cell type quantification in demyelinating lesions

Images were collected from 2-3 consecutive sections 144 µm apart containing the central part of the demyelinating lesion using a TCS STED CW SP8 super-resolution microscope (Leica) and imported into NIH ImageJ software. The area lacking MBP staining within the dorsal funiculus was delimited on a corresponding channel and imported onto the other channels to regionally restrict the analyses. For each image, the area of demyelination was measured and cells positive for the marker(s) of interest counted. Results are presented as the total number of positive cells per lesion area measured or the percentage of the total.

### Quantification of remyelination

Serial images were taken, in a blinded fashion, throughout the lesion to cover the demyelinated area using a transmission electron microscope JEOL JEM 1400 Plus (JEOL) connected to a digital camera (sCMOS), and imported into ImageJ. Remyelinated and demyelinated axons were counted, blindly. Results are presented as the percentage of the total axons counted.

### RNA extraction and microarray analyses

For each condition, we used four independent FACS samples, to provide four biological replicates. Total RNA was extracted using NucleoSpin RNA XS kit (Macherey-Nagel). Quantity and quality of RNA extractions were analyzed using Agilent RNA 6000 Pico kit (Agilent). Labeled RNAs (Liqa Kit, Agilent) were then hybridized onto Agilent whole-mouse genome microarray chips. Data were normalized and analyzed using the R statistical open tool (R Manuals; RRID:OMICS_01764). We used a Student’s t test and Benjamini–Hochberg test to identify the differentially expressed genes between two conditions (cutoffs: p < 0.001 and q < 0.01, respectively). A full list of genes was deposited in NCBI GEO (Gene Expression Omnibus; RRID:nif-0000-00142; accession number: GSE48872).

### RNA sequencing

For control and lesion oligodendroglial-enriched tissues (isolated following Percoll gradient cell separation from enzymatically dissociated spinal cord tissue)^20^, approximately 10 ng of total RNA per sample was used for library construction with the Ultra-Low-RNA-seq RNA Sample Prep Kit (Illumina) and sequenced using the Illumina HiSeq 4000 instrument according to the manufacturer’s instructions for 100bp paired end read runs. High-quality reads were aligned to the mouse reference genome (mm10), RefSeq exons, splicing junctions, and contamination databases (ribosome and mitochondria sequences) using the Burrows-Wheeler Aligner (BWA) algorithm. The read count for each RefSeq transcript was extracted using uniquely aligned reads to exon and splice-junction regions. The raw read counts were input into DESeq2 v1.2.5 81, for normalizing the signal for each transcript and to ascertain differential gene expression with associated p-values. We used a cut-off of p-value < 0.01 and q-value < 0.05 to identified differentially expressed genes. A full list of genes was deposited in NCBI GEO (Gene Expression Omnibus; RRID:nif-0000-00142; accession number: GSE122446).

### Statistics

All data are expressed as mean ± SEM. Statistical analyses were carried out using Prism software (GraphPad Software), with p values < 0.05 considered significant. First, data distribution was verified using Prism (D’Agostino and Pearson Normality test), and subsequently, the appropriate parametric versus non-parametric test was chosen. For IHC and EM analyses, as the normality test indicated, data were analyzed using non-parametric statistics, with Kruskal-Wallis test to compare 3 or more groups, followed by Tukey’s multiple comparison post-hoc test. Depending on the results of the Normality test, differences between 2 groups were analyzed using Mann-Whitney test or Unpaired student t-test (2-tailed). Numbers of replicates and p values relevant to specific data are presented in the figure legends.

### Data availability

Data are available upon reasonable request.

## RESULTS

### Donor-derived macrophages infiltrate demyelinating lesions

To investigate whether transplant-derived cells target demyelinating CNS lesions, we injected pre-conditioned mice with whole bone marrow (WBM) isolated from actin-GFP mice (Fig. 1A). At two months post-transplant, 72.6±5.8% of blood cells were GFP+. Moreover, 87.07±9.7% of blood CD11b+ cells (monocytes) were GFP+. Thus, monocytes in chimeric mice derive mostly from transplanted cells (Fig. 1B).

**Figure 1.**
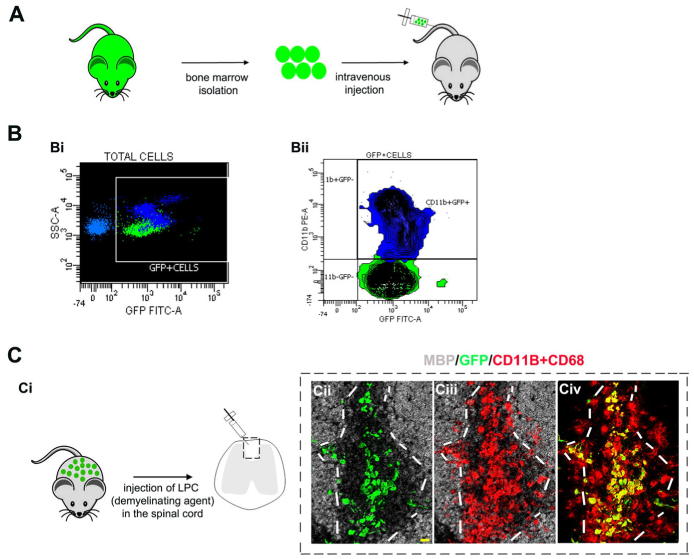
Graft-derived macrophages in demyelinating lesions. **A**. Bone marrow isolated from actin-GFP mice was transplanted into pre-conditioned recipients. **B**. Flow cytometry analyses of recipient blood 2 months post-transplant. Bi. Transplanted cells detected by GFP fluorescence. Bii. CD11b lebelling shows a large proportion of monocytes expressing GFP. **C**. Demyelinating lesions in chimeric mice at 3 dpl. Ci. Demyelination was induced by injecting LPC in the spinal cord white matter. Cii-Civ. Co-immunolabeling for GFP, MBP, and a mixture of CD11b/CD68, showing graft-derived macrophages in the lesion (identified by lack of MBP staining). White lines indicate lesion borders. Scale bar 20 µm. MBP-myelin basic protein; GFP-green fluorescent protein.

We next induced demyelination in the spinal cord of chimeric mice by injecting lysolecithin (LPC) (Fig. 1Ci).

At both 3 and 7 days post lesion (dpl) (ongoing OPC recruitment), GFP+ cells were found in demyelinated areas, and were CD11b/CD68+(macrophage marker) (Fig. 1Cii-Civ). Quantification revealed that 91.5±1.6% of GFP+ cells were macrophages. Moreover, 47.5±4.6% of macrophages in the lesion were GFP+. Because 96.35% (median) of circulating blood monocytes were GFP+, we assume that within lesions, all blood-derived cells are GFP+, while resident microglia/macrophages are GFP-, and thus conclude that both resident and blood-derived macrophages are recruited early to LPC lesions.

Therefore, graft-derived macrophages reliably target CNS lesions early after demyelination and hence represent appropriate molecular vehicles for stimulating OPC repopulation/myelin repair.

### Genetically-modified HSCs secrete Sema3F that increases OPC migration

Next, we generated lentiviral vectors to overexpress Sema3F. We used a bicistronic construction with self-cleaving T2A sequence inserted between Sema3F- and GFP-encoding sequences, all under PGK promoter (PGK-GFP-T2A-SemaF; Fig. S1). Thus, Sema3F and GFP should be produced as separate proteins.

Next, we genetically modified HSCs to overexpress Sema3F. HSCs were isolated, transduced with PGK-GFP-T2A-Sema3F or PGK-GFP lentiviral vectors, and expanded for 5 days (Fig. 2A). 35.8±5.3% of PGK-GFP-T2A-Sema3F- and 76.7±4.8% of PGK-GFP-transduced cells were GFP+ (Fig. 2B). This difference in transduction efficiency can be explained by a 30% greater size of Sema3F-GFP transgene (Fig. S1).

**Figure 2.**
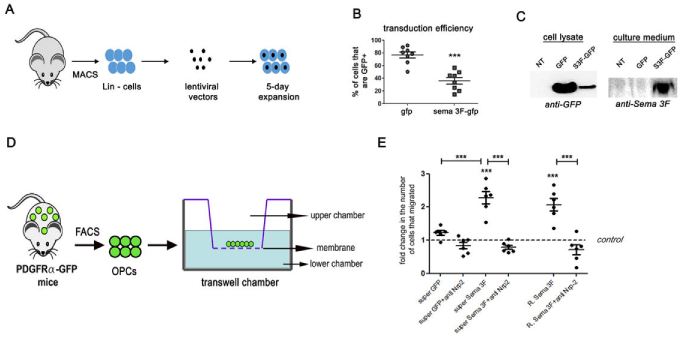
Genetically enginereed HSCs secrete Sema3F that attracts OPCs in vitro. **A-C**. Transduction of HSCs. **A**. Mouse bone marrow was isolated and differentiated cells (Lin+) were removed using MACS. Non-differentiated cells were transduced and expanded for 5 days. **B**. Lower numbers of GFP+ cells, verified using flow cytometry, after transduction with Sema3F vector, p<0.0001, Student t test, n=8 independent experiments. **C**. Western blot showing GFP expression in cells transduced with both vectors and Sema3F expression in the supernatant of Sema3F-transduced cells only. **D**. OPCs isolated from adult PDGFRα-GFP mice using FACS were subjected to transwell-chamber migration assay in response to supernatants from non-transduced (NT), GFP-transduced (gfp), or Sema3F -transduced (sema3F-gfp) HSCs. **E**. Fold increase in cells that crossed the membrane insert compared to NT. p<0.0001. One-way ANOVA followed by Bonferroni’s Multiple Comparison test. n=normalized replicates from 3-5 independent experiments. R.=recombinant. Super=supernatant. Anti-Nrp2=Neuropilin2 function blocking antibody.

Using western blot, we detected GFP expression in the lysates of both control and Sema3F-transduced cells, although to a lesser extent in the latter (Fig. 2C), consistent with lower transduction efficiency of PGK-GFP-T2A-Sema3F. Sema3F was detected exclusively in the supernatant of PGK-GFP-T2A-Sema3F-transduced cells (Fig. 2C).

Cell viability (propidium iodide and Viobility Dye labeling) and the pattern of colony formation (proliferation/differentiation potential) were similar between GFP- and Sema3F-transduced HSC preparations (Fig. S2).

To verify the functionality of Sema3F secreted by HSCs, we used transwell-chamber assay and quantified migration of adult OPCs in response to the supernatant from either non-transduced (NT)-, PGK-GFP- (GFP) or PGK-GFP-T2A-Sema3F-transduced (Sema3F-GFP) HSCs (Fig. 2D). Migration was 2-fold higher in Sema3F-GFP versus GFP (Fig. 2E), which was similar to the increase in migration induced by a recombinant Sema3F at 500 ng/mL (2.2 fold compared to control, Fig. 2E). An increase in OPC migration was also observed in experiments using supernatants from PGK-GFP-T2A-Sema3F or PGK-GFP transduced HEK cells (not shown).

The increase in migration induced by the supernatant from Sema3F-GFP-transduced HSCs was abrogated in the presence of anti-Nrp2 function blocking antibody. Similar findings were obtained for recombinant Sema3F (Fig. 2E). Thus, Sema3F-GFP transduced HSC supernatant increases OPC migration in vitro in Nrp2-dependent manner.

Therefore, while transduction with PGK-GFP-T2A-Sema3F does not affect HSC viability or differentiation in vitro, it leads to secretion of Sema3F that increases OPC migration, via Nrp2.

### OPCs from both young and medium-age mice are attracted by Sema3F

Next, we investigated whether the responsiveness of OPCs to Sema3F decreases with age. We first checked whether aged OPCs retain Nrp2 expression. Both microarray data on FACS-sorted OPCs from the brains of PDGFRα-GFP mice (Fig. 3A) and RNA sequencing data on oligodendroglial spinal cord cells (Fig. 3B) demonstrated lack of statistically significant changes in Nrp2 mRNA expression between 2 month-old (young) and 12 month-old (middle-aged) mice. Interestingly, immunohistochemical analyses of Sema3F expression in LPC lesions at 4 dpl (Fig. 3C-E) and 7 dpl showed a significant decrease in Sema3F-expressing cells in lesions induced in middle-aged and old (18 month-old) mice at 4 dpl (onset of OPC recruitment), as compared to young mice (Fig. 3F), which was partially due to a decrease in microglial/macrophage Sema3F expression (Fig. 3G). We then evaluated whether recombinant Sema3F or the supernatant from PGK-GFP-T2A-Sema3F-transduced cells retains its capacity to attract OPCs with age. We exposed OPCs FACS-sorted from the brains of young versus middle-aged PDGFRα-GFP mice to recombinant Sema3F or PGK-GFP-T2A-Sema3F-transduced cell culture supernatant in a Boyden transwell chamber for 24 h. Our results show that OPCs isolated from middle-aged mice are mobilized by both recombinant Sema3F and supernatant-derived PGK-GFP-T2A-Sema3F-transduced cell supernatant (Fig. 3H). We have not performed migration experiments with OPCs isolated from old (18m of age) mice because, using FACS sorting, we obtain very low numbers of cells, insufficient for migration experiments.

**Figure 3.**
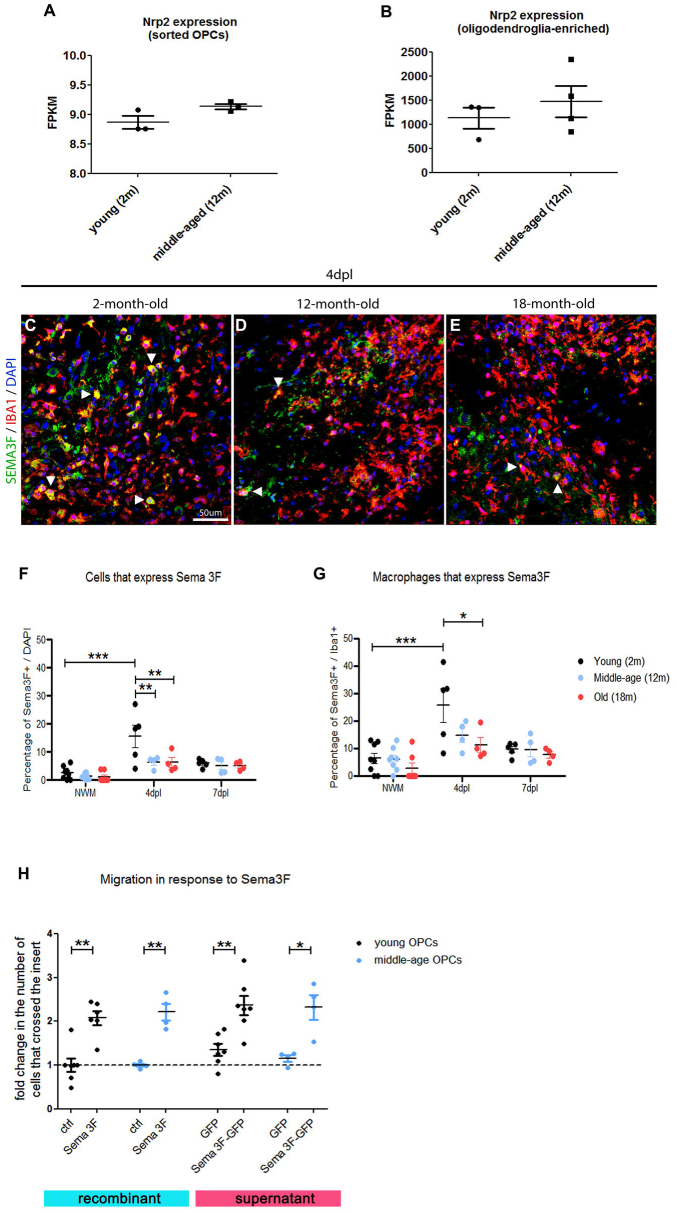
Middle-age OPCs maintain Nrp2 expression and responsiveness to Sema3F. **A-B.** Expression of Nrp2 gene by OPCs is maintained in middle-age (12 month (m) old) mice. A. Microarray analyses results for Nrp2 expression by OPCs purified using FACS from the brains of 2 months versus 12 months old PDGFRa −GFP mice (n=3-4 independent experiments). B. RNA sequencing results for Nrp2 expression by oligodendroglia-enriched preparations isolated from the spinal cords of 2 month versus 12 month old mice. (n=3-4 independent experiments). **C-G**. Sema 3F expression in demyelinating lesions at 4 dpl (onset of OPC recruitment) is decreased in middle-aged (12 m old) and old (18m old) mice compared to young mice (2m old). C-E. Colabellings for Sema3F and Iba1 (macrophage marker) in demyelinating lesions at 4 dpl in 2m (C), 12m (D),and 18m old mice (E). Arrowheads point to double-labelled cells. F. Quantification of Sema3F expressing cells in the lesion. G. Percentage of Iba1+ cells positive for Sema3F. NWM-normal white matter. dpl-days post lesion. n=4-8 mice/group. Two-way ANOVA followed by Tukey’s Multiple Comparison test. **H**. Transwell chamber assay was used to evaluate migration of young versus middle-age OPCs in response to recombinant Sema3F and supernatant from PGK-GFP-T2A-Sema3F transduced cells. OPCs of both ages migrate in response to recombinant Sema3F and PGK-GFP-T2A-Sema3F-transduced cell supernatant. n=normalized replicates from 2-5 independent experiments. One-way ANOVA followed by Bonferroni’s Multiple Comparison test. *p<0.05, ** p<0.01. Scale bar 50 µm.

Thus, OPCs from middle-aged mice retain the expression of Nrp2 gene and are attracted by Sema3F.

### Hematopoietic reconstitution by transduced HSCs

We transplanted female GFP and Sema3F-transduced HSCs (mixture of transduced and non-transduced cells) to preconditioned male mice (GFP and Sema3F mice; Fig. 4A). At 2 months post-grafting, qPCR for sex-containing region of the Y chromosome showed extensive reconstitution by transplanted (female) cells in both groups (87.96±4.05% GFP and 88.68±2.41% Sema3F) (Fig. 4B). Flow cytometry analyses of GFP expression, revealed 73.8±3.1% of GFP+ (transgene-carrying) cells in GFP and 22.9±2.1% in Sema3F group (Fig. 4C), consistent with lower transduction efficiency of PGK-GFP-T2A-Sema3F vector mentioned above.

**Figure 4.**
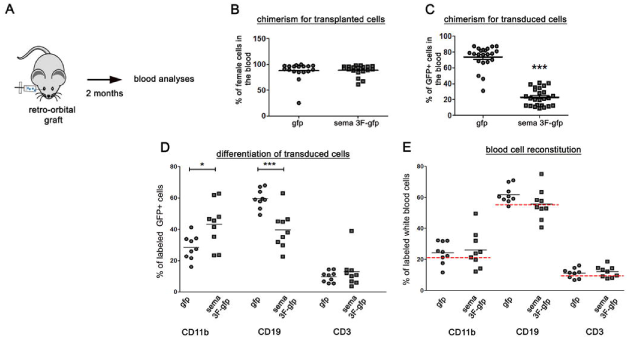
Generation of Sema3F chimeras. **A**. Transduced female HSCs were injected retro-orbitally into pre-conditioned male mice. Blood was analyzed 2 months post-transplant. **B**. Percentage of transplanted (female) cells in the blood. n=18-19 mice/group. **C**. Percentage of transduced (GFP+) cells in the blood. p<0.0001. n=22-25 mice/group. **D**. Percentages of GFP+ cells labeled with CD11b, CD19, and CD3. The proportion of GFP+ cells labeled with CD11b is increased in Sema3F mice (p=0.01), while that of CD19-labeled cells is decreased (p=0.0003). Unpaired Student t-test. **E**. Percentages of total blood cells expressing CD11b, CD19, and CD3. Red dashed lines indicate the percentage expression of these antigens in blood cells of control mice. n=9 mice/group.

Using flow cytometry, we analyzed differentiation of transduced (GFP+) cells into monocyte (CD11b+), B cell (CD19+), and T cell (CD3+) lineages. Interestingly, the percentage of GFP cells co-expressing CD11b was higher in Sema3F mice, while that co-expressing CD19 was lower (Fig. 4D). However, this difference did not affect the blood cell composition in chimeric mice as the percentages of total CD11b+, CD19+, and CD3+ cells in the blood were similar between the two groups and comparable to that of control (non-treated) mice (Fig. 4E). Therefore, Sema3F mice show normal blood composition.

We then analyzed whether Sema3F modifies GFP+ cell numbers in the heart, liver, lung and spleen. We detected numerous GFP+ cells in the spleen, as expected. We also detected GFP+ cells in the lungs, albeit at lower numbers as compared to the spleen. Correlation analyses revealed that these numbers were determined by the GFP chimerism in the blood (r = 0.84 spleen, r = 0.94 lung), which indicated that Sema3F expression did not alter GFP+ cell homing and maintenance in these organs.

### Sema3F-carrying cells successfully target demyelinating lesions and are predominantly macrophages

We investigated the response of transgene-carrying cells to LPC-induced spinal cord demyelination. GFP+ cells were detected at 7 and 10 dpl in GFP and Sema3F mice with less cells in the latter, although this difference was not statistically significant (Fig. 5A-B, D-E, G). Using Prism, we detected a strong correlation between the percentage of GFP+ cells in the blood and numbers of GFP+ cells in lesions at both 7 (r =0.88) and 10 dpl (r =0.81). This indicates that numbers of GFP+ cells in the lesions is determined by GFP blood chimerism, itself dependent on transduction efficiency (lower for PGK-GFP-T2A-Sema3F). Therefore, Sema3F expression does not alter monocyte homing to demyelinating CNS lesions.

**Figure 5.**
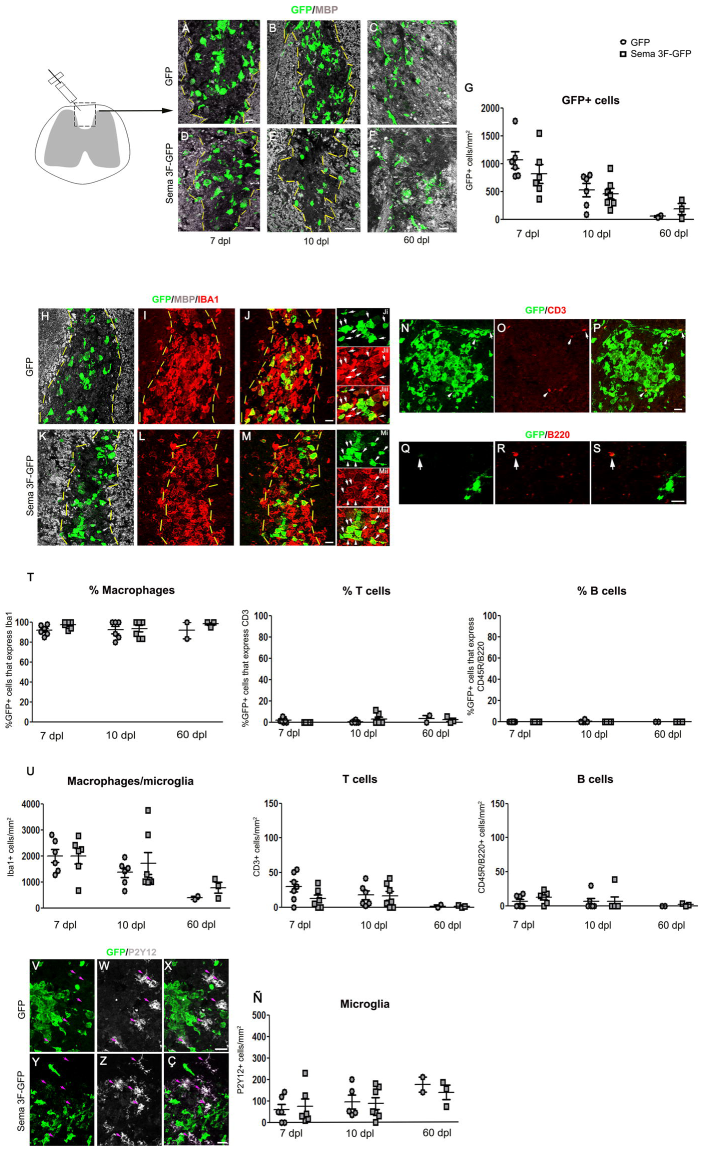
Transgene-carrying macrophages in demyelinating lesions. **A-F.** Co-labeling for GFP and MBP on spinal cord sections of GFP (A-C) and Sema3F-GFP chimeras (D-F) sacrificed at 7 (A,D), 10 (B,E), and 60 (C,F) days post lesion (dpl). Absence of MBP labeling in A-B,D-E indicates the lesion (delimited by yellow lines). Areas faintly labeled with MBP in C-F likely represent remyelinated areas. **G**. Quantification of GFP+ cells. Kruskal-Wallis test p=0.02. No significant differences between GFP and Sema3F at any of the time points (Dunn’s post test; Mann-Whitney test). Significant decrease in GFP+ cells between 7 and 60 dpl for both GFP and Sema3F mice 2m (Dunn’s post test). **H-M**. Co-labeling for GFP/MBP/Iba1 (macrophage/microglia marker) shows colocalization of GFP with Iba1. **N-P** Co-labeling for GFP and T cell marker CD3. **Q-S**. Co-labeling for GFP and B cell marker CD45R/B220. **T**. Percentages of GFP+ cells that are Iba1+, CD3+, and CD45R/B220+. The absolute majority of GFP+ cells are Iba1+. **U**. Quantification of Iba1+, CD3+, and B220+ cells. For all 3 populations, no significant differences in cell numbers were observed between GFP and Sema3F mice, and a general decrease was observed at 60 dpl compared to earlier time points. **V-Ç**. Co-labeling for GFP and microglial marker P2Y12. Purple arrows indicate P2Y12+ cells. Exclusion between the two markers is obvious in both groups. **Ñ**. Quantification of P2Y12+ cells. No significant changes are observed between the groups or across the time. For all figure panels, n=6-7 mice/group for 7 and 10 dpl, n=2-3 mice/group for 60 dpl. Scale bars 20 µm, except for S (10 µm).

We analyzed whether recruited cells persist in the lesions at late time points. At 60 dpl, GFP+ cell numbers decreased compared to earlier time points in both GFP and Sema3F mice (GFP mice: Kruskal-Wallis test for GFP+ cells at 7,10, 60 dpl: p=0.0079; significant difference between 7 and 60 dpl, Dunn’s multiple comparison test; Sema 3F-GFP mice: Kruskal-Wallis test at 7,10, 60 dpl: p=0.0252; significant difference between 7 and 60 dpl, Dunn’s multiple comparison test). There were no significant differences in the numbers of GFP+ cells between the two groups at any of the time points analyzed (Fig. 5C, F-G). However, we could only analyze two GFP and three Sema3F chimeras, so we cannot deduce whether Sema3F-expressing cells are differentially retained in the CNS.

Co-immunolabeling with GFP and microglia/macrophage marker Iba1, T-cell marker CD3, and B-cell marker CD45R/B220 revealed that the absolute majority of GFP+ cells were macrophages (median range across all groups tested 91.67-100%) (Fig. 5H-M, T). Total Iba1+ cell numbers (infiltrating and resident macrophages + microglia) were similar in GFP versus Sema3F mice (Fig. 5U), inferring that total myeloid infiltration was unchanged between these two groups. These numbers progressively decreased in time, as expected. Importantly, myelin debris removal (performed by macrophages/microglia) did not appear different between GFP and Sema3F chimeras, as evidenced by Oil Red O staining at 7 dpl (Appendix, Fig. S3).

Small numbers of GFP+ T cells were also detected in the lesions of both GFP and Sema3F chimeras (Fig. 5N-P, T, U). B cells were observed mostly in the meninges, in low numbers, and very few, that were GFP-, were also found in the spinal cord parenchyma at 7 dpl (Fig. 5Q-S, T, U). No changes were detected in the total numbers of infiltrating CD3+ and B220/CD45+ cells between GFP and Sema3F chimeras at any of the time points analyzed (Fig. 5T).

We then investigated whether Sema3F overexpression by infiltrating cells altered numbers of microglial cells in lesioned areas by performing immunohistochemistry for microglial marker P2Y12. In both groups, we observed a clear distinction between GFP+ and P2Y12+ cells (Fig.5V-Ç), the latter observed preferentially around lesion borders and outside the lesion. Unlike the numbers of GFP+ cells or total Iba1+ cells, numbers of P2Y12+ cells did not decrease with time (Fig. 5Ñ). However, it is possible that not all microglial cells were detected by P2Y12 labeling because, under certain pathological circumstances, these downregulate markers such as P2Y12 and TMEM119^21^.

Thus, the absolute majority of transgene-carrying cells in LPC lesions are macrophages, which is consistent with previous reports showing scarce lymphocytic infiltration in toxin-induced demyelinating lesions^22–24^. In addition, the inflammatory status of LPC lesions in Sema3F chimeras is comparable to that of the control mice.

Interestingly, while basically all GFP+ cells at 7 and 10 dpl had morphology of myelin-phagocyting macrophages (large, round), we also observed ramified GFP+ cells in the lesion-neighboring tissue at all time points and within the lesion at 60 dpl (Appendix, Fig. S4). This suggests that blood-derived myeloid cells 1) adopt microglia-like morphology in response to CNS milieu and 2) that a small portion of infiltrating cells remains in the CNS once lesions have remyelinated.

### Lesions in Sema3F mice show increased OPC recruitment

We then investigated whether Sema3F overexpression by blood-derived cells in demyelinating lesions affects OPC repopulation of the lesion. Numbers of the PDGFRα+ (OPCs) and Olig2+ (oligodendroglia) cells were quantified at 7 and 10 dpl, as OPC recruitment/proliferation is maximal within this time window (Fig. 6A), and were significantly higher in Sema3F mice at 7 dpl (Fig. 6B-D, E-G). We investigated whether increased OPC numbers correlated with increased OPC proliferation in the lesions. Co-immunolabeling for Nkx2.2 (marker of reactive OPCs) and proliferation marker Ki67 showed that numbers of total Nkx2.2+ cells (Fig. 6H-I, J) were significantly higher in the lesions of Sema3F mice as compared to GFP mice. Nkx2.2 cells were observed in the proximity of GFP+ cells in the Sema3F group (Fig 6I). An increase in Nkx2.2+Ki67+ cells was also detected, although it was not statistically significant due to an outlier in the GFP group (Grubb’s test)(Fig. 6K-P, Q). Yet, the proportion of Nkx2.2 cells that was expressing Ki67 was similar between the two groups (Fig. 6R). These results suggest that Sema3F lesions recruit more OPCs, rather than significantly modifying their proliferation rate.

**Figure 6.**
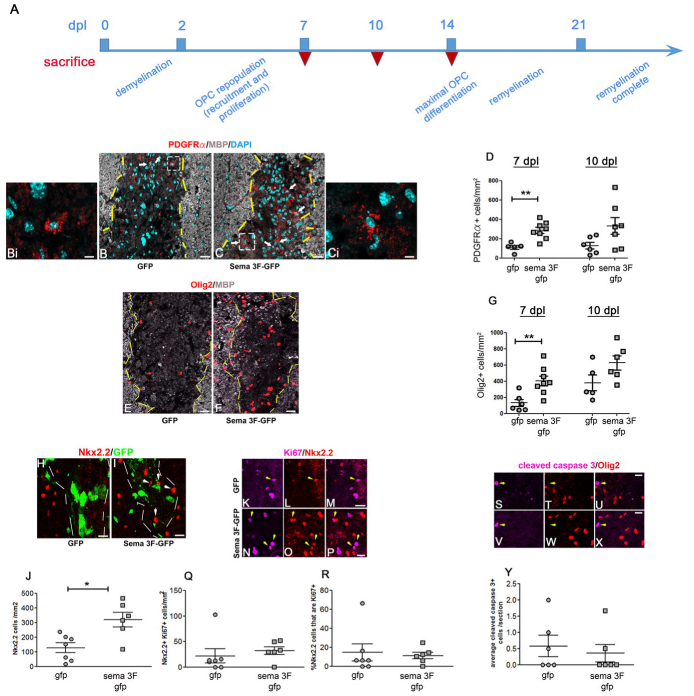
OPC recruitment is increased in Sema3F chimeras. **A**. Temporal sequence of events in LPC lesion. dpl-days post lesion. Red inverted triangles indicate sacrifice. **B-C, E-F**. Lesions at 7 dpl in GFP (B,E) and Sema3F (C,F) mice. B-C. Co-labeling for PDGFRα, MBP, and DAPI. Arrows indicate OPCs in the lesion. Bi and Ci-insets of white dotted squares in B and C. **D**. Higher numbers of PDGFRα+ (OPCs) cells in Sema3F mice at 7 dpl (Mann Whitney test: p=0.0031, n=6-8 mice/group). E-F. Co-labeling for Olig2 and MBP. **G**. Higher numbers of Olig2+ (oligodendroglial) cells in Sema3F-GFP mice at 7 dpl (Mann Whitney test: p=0.0047, n=5-8 mice/group). **H-I**. Immunolabelling for activated OPC marker Nkx2.2. **J**. Numbers of Nkx2.2+ cells are increased in lesions of Sema3F-GFP mice at 7dpl (Mann Whitney test: p=0.014, n=6-7 mice/group). **K-P**. colabelling for Nkx2.2 and proliferation marker Ki67. **Q**. Numbers of Nkx2.2+Ki67+ cells are increased at 7 dpl in Sema3F-GFP mice. **R**. The proportion of Nkx2.2+ cells co-expressing Ki67 is unchanged. **S-X**. Colabelling for Olig2 and cleaved caspase 3, marker of apoptosis indicates occasional apoptotic cells in both groups of mice that are Olig2-. **Y**. Numbers of cleaved caspase 3+ cells are the same in two groups. Scale bars 20 µm (B,C,E,F), 10 µm (H,I,M,P,U,X), 5 µm (Bi, Ci).

We also investigated whether oligodendroglial apoptosis is modified in Sema3F mice. Co-labeling for Olig2 (marker of OPCs and oligodendrocytes) and apoptosis marker cleaved caspase 3 revealed low numbers of apoptotic cells in both GFP and Sema3F mice lesion/perilesion that were Olig2- (Fig. 6S-Y). Thus, presence of Sema3F-expressing blood derived cells does not affect oligodendroglial apoptosis.

### Increased oligodendrogenesis and remyelination in Sema3F mice

New oligodendrocytes in LPC lesions appear during second week post LPC (Fig. 6A). Immunolabelings for APC/CC1+ cells (oligodendrocytes) in the lesions of GFP and Sema 3F mice at 7 and 10 dpl showed increased numbers of these cells in Sema3F mice at both time points (Fig. 7A-C). Thus, increased OPC recruitment in Sema3F mice is followed by accelerated generation of new oligodendrocytes.

**Figure 7.**
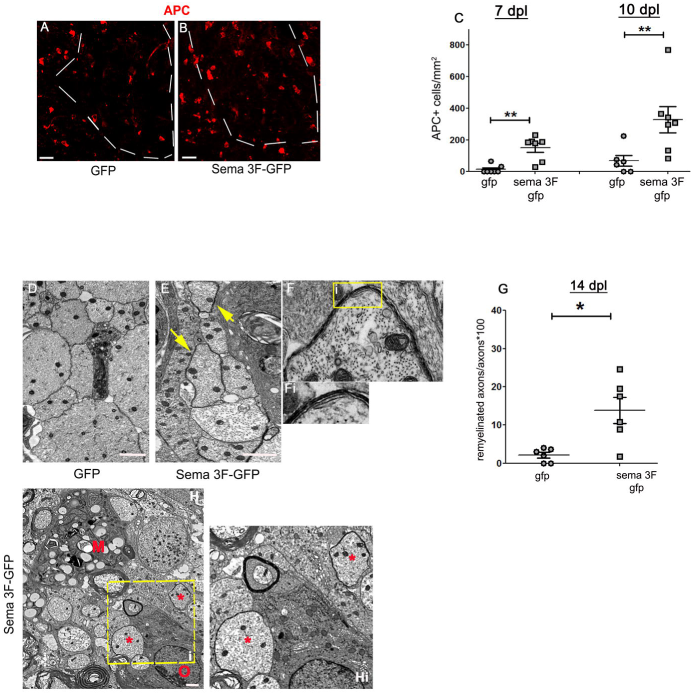
Oligodendrogenesis and remyelination are accelerated in Sema 3F chimeras. **A-B**. Labelling for APC/CC1, a marker of oligodendrocytes. Dotted lines indicate the lesion border. APC+ cells are scarce in the lesions of GFP mice (A). B. APC+ cells in the lesion of Sema3F mice. **C**. Quantification of APC+ cells (Mann Whitney test; p=0.0061 for 7 dpl, p=0.0081 for 10 dpl; n= 6-7 mice/group). **D-E**. EM images of the lesion in GFP (D) and Sema3F (E) chimeras at 14 dpl. Thin myelin sheaths in Sema 3F mice (yellow arrows). **F**. Higher power image of a new myelin sheath in a Sema3F mouse. **Fi**. Inset of yellow square in F. **G**. Quantification of remyelinated axons (Mann Whitney test: p=0.03, n=6 mice/group). **H.** In Sema3F mice, oligodendroglial cells (O) and early remyelination (*) are frequently observed in macrophage (M) vicinity. **Hi**. Inset of yellow dashed square in H. Scale bars 20 µm (A,B), 1 µm (D,E).

We then analyzed whether accelerated oligodendrogenesis also accelerates myelin repair. We used transmission electron microscopy (TEM) to quantify axons without myelin and those surrounded by thin myelin sheaths in the lesions at the onset of remyelination (14 dpl). While at this early time point demyelinated axons were predominant in both groups, the proportion of axons surrounded by thin myelin sheaths was higher in Sema3F mice (2.12±0.73% in GFP versus 13.83±3.39% in Sema3F mice) (Fig. 7D-G). The axons undergoing remyelination were frequently seen in the vicinity of macrophages (Fig. 7H). Thus, Sema3F mice show a faster onset of remyelination.

Therefore, recruitment of Sema3F-expressing blood-derived macrophages to focal demyelinating lesions increases OPC repopulation, which accelerates oligodendrogenesis and remyelination.

## DISCUSSION

Here, we have described a clinically-relevant paradigm for targeting Sema3F to demyelinating lesions. Using transplantation of genetically-modified HSCs we showed that Sema3F-carrying blood-derived macrophages target areas of demyelination very quickly after demyelination, which results in increased OPC recruitment and accelerated remyelination. Increasing OPC recruitment and accelerating remyelination is likely to benefit patients with MS for several reasons:

1. Acute inflammation in demyelinating lesions is beneficial and even necessary for successful myelin repair^25,26^. This means that quick recruitment of OPCs to areas of demyelination before the complete resolution of inflammation will enhance repair by these cells.
2. With time, axons undergo changes that may render them refractory to remyelination^27^, which means that the more quickly the OPCs interact with the demyelinated axons the more likely is that remyelination will take place.
3. Within the (likely short) window of time when axonal pathology is still reversible, remyelination protects axons from degeneration^2^.
4. As previously shown, most OPCs in early MS lesions undergo differentiation and very few proliferate^6,7,28^, which suggests that repeated rounds of demyelination observed in MS^29,30^ may exhaust the OPC pool and lead to remyelination failure^6^. Thus, replenishing the OPC pool during demyelination should benefit remyelination and enhance axonal protection in long term, thus potentially preventing/significantly delaying lesion chronicity and the development of permanent neurological deficit.

We used a lysolecithin model to investigate whether OPC recruitment and remyelination can be enhanced by targeting Sema3F expression to demyelinating lesions via blood-born macrophages. The advantage of this model of acute demyelination induced by focal injection of membrane/myelin toxin lysolecithin is that the induction of demyelination is largely independent from blood cells. This allowed us to study the effect of Sema3F overexpression in monocytes/macrophages on myelin repair without confounding effects on the extent of demyelination. While it would have been interesting to confirm our results in a model of diffuse demyelination (cuprizone intoxication), or autoimmune CNS inflammation (experimental autoimmune encephalomyelitis, EAE), the severity and high mortality risk in these models, incompatible with preconditioning protocols, precluded us from performing these experiments.

Because of the nature of the lysolecithin model (with extensive myeloid infiltration and scarce presence of other immune cell types), the absolute majority of the infiltrating cells observed in our experiments were macrophages. Presence of Sema3F-expressing macrophages in the lesions did not affect the overall inflammatory status of the lesions as total numbers of macrophages, T cells, B cells, and microglia remained unchanged. It did not affect apoptosis in the lesioned tissue either.

Thus, given that neither inflammatory status of the lesion nor OPC survival were affected in Sema3F chimeras and that Nrp2 (Sema3F receptor) function-blocking antibody abrogated OPC migration in response to Sema3F-transduced HSC supernatant, we conclude that increased OPC recruitment in Sema3F chimeras is mediated by the interaction of blood cell-secreted Sema3F and Nrp2 expressed on OPCs.

Progression in MS tends to start in the early fifth decade of life (middle-age)^31^. Importantly, in animal models, the efficiency of both OPC recruitment and remyelination decreases with age^32^, which is already evident in middle-aged mice (9-12 months old). We investigated whether Nrp2 expression in OPCs diminishes with age thus decreasing their responsiveness to Sema3F. Interestingly, we found that Nrp2 expression on OPCs is maintained in middle-aged mice, but the number of Sema3F-expressing cells in lesions is significantly lower in middle-aged and old animals. This suggests that OPC recruitment in older animals may be impaired, at least in part, due to decreased expression of Sema3F in lesions rather than OPC unresponsiveness to this molecule. Importantly, we showed that OPCs isolated from middle-aged animals efficiently migrate in response to both recombinant Sema3F and the supernatant from Sema3F-transduced HSCs. While we do think that overexpressing Sema3F in MS lesions as early as possible (young patients with relapsing-remitting disease) would be the most optimal in stimulating remyelination to prevent neurodegeneration, conserved OPC responses to Sema3F in middle-aged mice suggest that such therapy may also be beneficial in early stages of progressive disease (most frequently occurring in middle-age individuals) to limit further axonal loss by enhancing remyelination of expanding lesions.

OPC repopulation of the lesions was enhanced in Sema3F chimeras at 7 dpl. Even though such repopulation is dependent on both OPC recruitment and proliferation, we think that Sema3F predominantly increased the recruitment because of the following:

1. As we showed previously^10^ and in this article, in vitro exposure of adult OPCs to Sema3F increases OPC migration, but has little effect on proliferation.
2. The proportion of OPCs undergoing proliferation was similar in control and Sema3F chimeras at 7 dpl (proliferation peak).

Most importantly, increased numbers of OPCs during recruitment phase, achieved using hematopoietic cell gene therapy, were associated with accelerated myelin repair. We think these findings have important implications for the development of remyelination-promoting therapies in MS. In addition to promoting OPC recruitment, which, if performed during early phases of the disease could promote remyelination and neuroprotection thus preventing disease chronicity, this paradigm could also be used to overexpress neurotrophic factors and anti-oxidant molecules, potentially beneficial for progressive MS^33^.

Developing strategies to prevent, as well as to treat progressive MS is an unmet therapeutic need. We believe that successful remyelination-promoting strategies in relapsing-remitting MS should greatly reduce the likelihood of progressive disease, and efficient targeting of repair-promoting molecule appears crucial to achieve this. The strategy we are proposing relies on blood-derived macrophage infiltration in CNS lesions that are undergoing activity, both in RRMS and early progressive MS, while axons are still viable. Interestingly, the treatment with busulfan, the drug used as a pre-conditioning treatment for HSCT, enhances CNS engraftment of blood monocytes derived from grafted HSCs in the absence of blood-brain barrier breakdown^34,35^. This explains the benefits of genetically-modified HSCT described in patients with leukodystrophies, and suggests that MS lesions in the progressive phase (with no breakdown of blood-brain barrier) may also be targeted by transgene-carrying blood monocytes to enhance remyelination. In addition, as mentioned above, macrophage-mediated delivery of neurotrophic factors to the CNS may be beneficial in preventing/reducing global CNS neurodegeneration observed in progressive disease.

We have used transplantation of HSCs as means of obtaining transgene-carrying monocytes/macrophages. Autologous HSC transplantation (aHSCT) has been used to stabilize the disease in a defined population of patients with RRMS, with little or no benefits observed in progressive disease^36^, likely because suppressing inflammation is not sufficient to promote remyelination at this stage. Theoretically, combining genetic modification of HSCs with autologous transplantation in clinic might potentially both regulate inflammation and promote regeneration, thus extending the benefits of aHSCT to a wider subset of eligible patients. However, it must be taken into account that aHSCT is associated with significant side effects, in part due to preconditioning regimens, and is therefore recommended only for patients with very aggressive inflammatory disease and resistant to conventional therapies^36^. It is important to highlight that application of genetically-modified macrophages to increase remyelination could also be achieved using alternative strategies. One of these, potentially applicable to a larger population of patients (in which aHSCT is not recommended), may be to systemically deliver engineered monocytes amplified from autologous progenitor cells, as previously described^37^, given that demyelinating MS lesions are always associated with the presence of myeloid-lineage cells^1^. Future work should test this strategy, that does not require pre-conditioning, first in rodent models such as the lysolecithin injection, cuprizone intoxication, and EAE, and then, if benefits are confirmed, in marmoset EAE, considered as the most suitable pre-clinical model of MS^38,39^. Moreover, it may be possible to restrict the expression of the transgene using specific promoters, so that the expression becomes upregulated specifically when the cell is performing phagocytosis (myelin debris), and thus reduce potential side effects in non-lesioned tissue, even though this may not be crucial if one aims to overexpress neurotrophins/anti-oxidants.

In conclusion, we provide a proof-of-concept that targeted molecular delivery by genetically-modified blood-derived macrophages increases OPC recruitment and accelerates remyelination. Although further data are required to clearly define the suitability of this strategy for specific patient subpopulations, these results and recent advances in gene therapy for human CNS diseases^40^ suggest our approach as a potentially useful therapeutic strategy to prevent/improve disability in MS.

## Author contributions

Conceived project VT, NC, and CL; Performed experiments: MSA, GDC, SM, BGL, ST, LI, CB, LBC, JCC, VT; Analyzed data: MSA, GDC, SM, BGL, ST, VT; Manuscript writing: VT with input from NC and CL. Assured funding: CL, NC, VT, and CM.

## Acknowledgements

We thank Dominique Langui and Asha Baskaran (BioImaging Facility-ICMQuant), Catherine Blanc and Benedicte Hoareau (Flow Cytometry Core CyPS, Pitié-Salpêtrière Hospital), Laura Escobar (Achucarro Basque Center for Neuroscience), and Ricardo Andrade (Microscopy Platform UPV/EHU) for technical assistance; Alexis Bemelmans and Noelle Dufour (Vector production platform, MIRCEN, CEA,) and Philippe Ravassard (ICM) for lentiviral vector production. We also thank Carlos Belmonte, Maria Domercq, and Sergio Lopez for comments on the manuscript. This study was performed at the cell culture (Celis) and preclinical functional exploration platform (PHENO-ICMice) core facilities of the ICM (Paris, France), Achucarro Basque Center for Neuroscience (Leioa, Spain), “Analytical and High-resolution Microscopy in Biomedicine Platform” of the University of the Basque Country (Leioa, Spain), and CUNY (NY, USA).

## Funding

This work was supported by the French National Institute of Health and Medical Research (INSERM), French National Research Agency (ANR, project Stemimus ANR-12-BSV4-0002-02), the European Leukodystrophy Association (ELA, project 2016-004C5B), NeurATRIS, the program “Investissements d’avenir” (ANR-10-IAIHU-06), CIBERNED (CB06/0005/0076), and Gobierno Vasco (IT1203-19). VT was a recipient of the Spanish Ministry of Economy Young Investigator Grant (SAF2015-74332-JIN).

## Conflict of interest

NC-Employed by Asklepios Biopharmaceutics. Collaboration with Bluebird Bio; CL-participation to advisory boards for Roche, Biogen, Merck-Serono, Genzyme, Vertex, Rewind; scientific collaboration with Vertex and Merck-Serono.

## Notes

### Summary of Updates

We have increased animal numbers in our analyses of OPC recruitment differentiation, and remyelination and added new data in the text, in the Figure 5, and supplementary figures. We have also modified the Discussion

