## Supplemental figures for "Genetically-Modified Macrophages Accelerate Myelin Repair"

### APPENDIX

**Table of content:**

**Figure S1 and legend:** Scheme illustrating lentiviral vector constructions used in the study

**Figure S2 and legend:** Viability and proliferation/differentiation following Sema3F transduction in vitro

**Figure S3 and legend:** Oil Red O staining in chimeric mouse spinal cord sections at 7 days post lesion

**Figure S4 and legend:** Morphology of GFP+ cells in the spinal cord

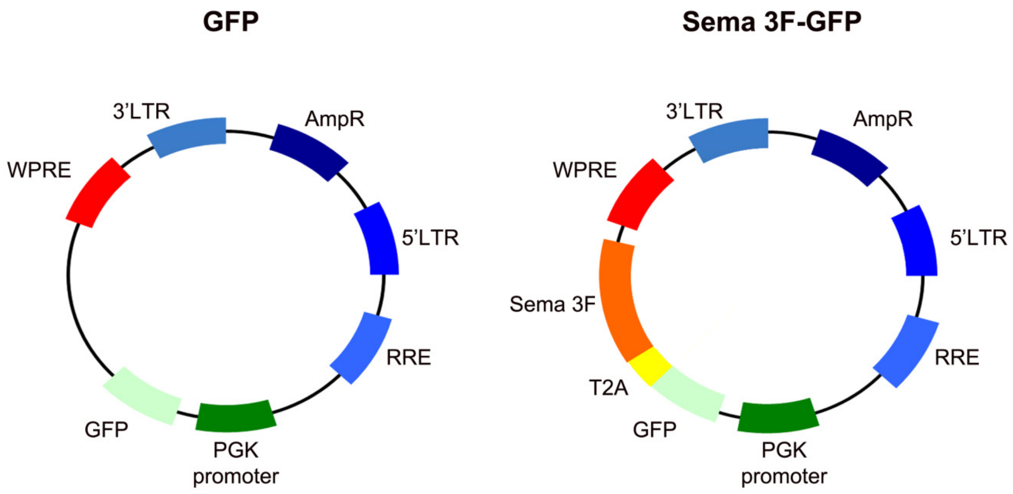


**Figure S1. Scheme illustrating lentiviral vector constructions used in the study.** WPRE-Woodchuck hepatitis virus Posttranscriptional Regulatory Element; LTR-Long Terminal Repeat; AmpR-Ampicillin Resistance; RRE- Rev Response Element; PGK-PhosphoGlycerate Kinase; GFP-Green Fluorescent protein; Sema-Semaphorin.

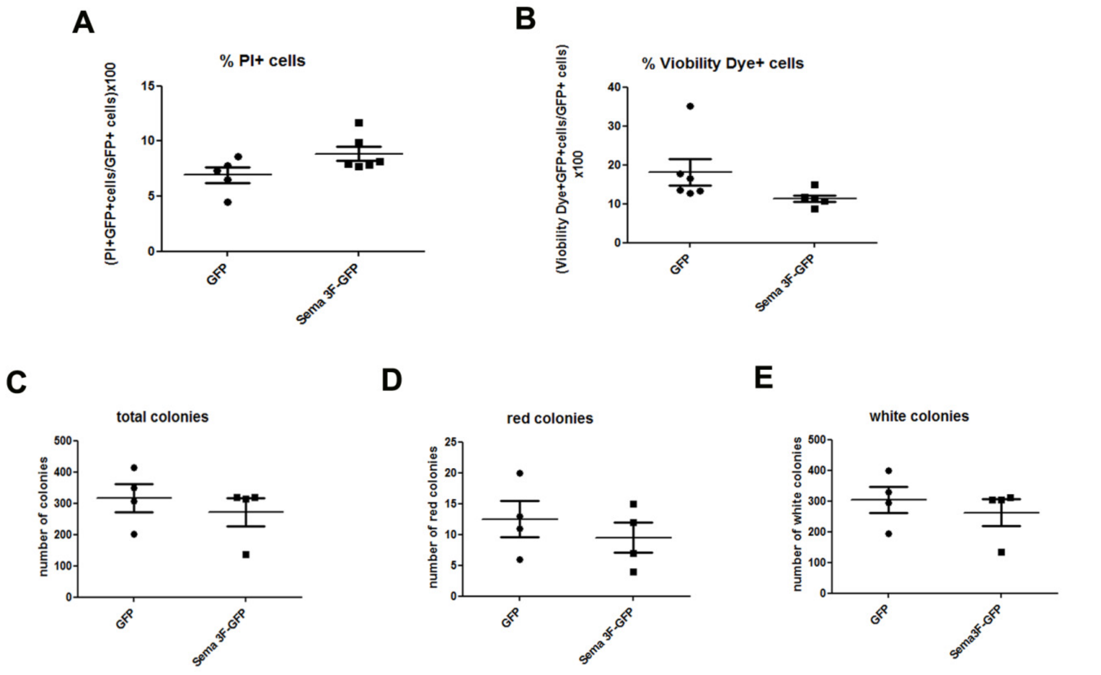


**Fig S2. Viability and proliferation/differentiation following Sema3F transduction in vitro. A-B**. Flow cytometry analyses of cell death. Hematopoietic stem/progenitor cell preparation was transduced in vitro using PGK-GFP and PGK-Sema3F-GFP lentiviral vectors and expanded for 5 days. Dead/apoptotic cells were labeled using Propidium Iodide (PI) and Viobility 405/520 Fixable Dye. n= 5-6 independent experiments. **A**. PI labelling. **B**. Viobility Dye labelling. **C-E**. Colony assay. n = replicates from 2 independent experiments. C. Total colonies formed by transduced cell preparations. D. Red colonies formed by transduced cell preparations. E. White colonies formed by transduced cell preparations.

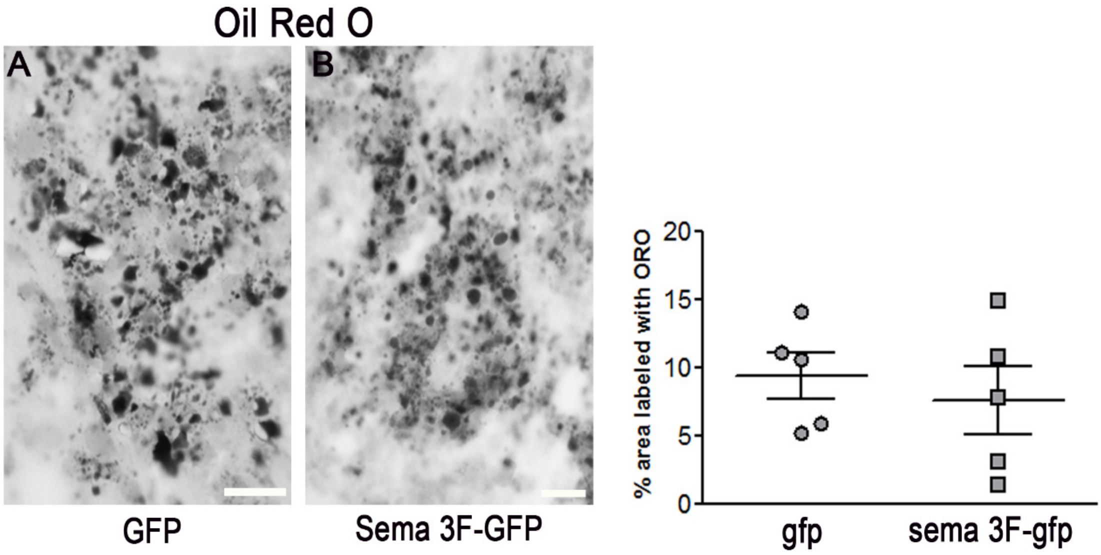


**Figure S3. Oil Red O staining in chimeric mouse spinal cord sections at 7 days post lesion.** **A**. GFP mouse. **B**. Sema3F-GFP mouse. C. The extent of labelling is similar between the two groups. n = 5 mice/group. Scale bars = 50 µm.

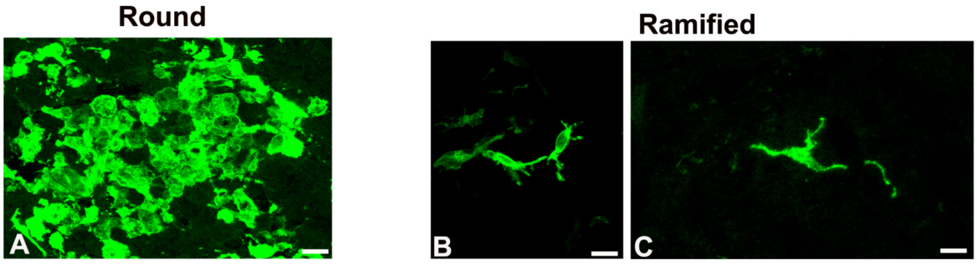


**Figure S4. Morphology of GFP+ cells in the spinal cord. A**. Large, round cells observed in the demyelinating lesions at all time points. This particular image is of a lesion at 7 dpl. **B-C**. Ramified cells observed in the lesion-neighbouring tissue at 7 and 10 dpl, but also in the lesions at 60 dpl. Scale bars A=20 µm, B=10 µm, and C=10 µm.
